# Tuning Cell-free Composition Controls the Time-delay, Dynamics, and Productivity of TX-TL Expression

**DOI:** 10.1101/2021.04.02.438196

**Authors:** Grace E. Vezeau, Howard M. Salis

## Abstract

The composition of TX-TL cell-free expression systems are adjusted by adding macromolecular crowding agents and salts. However, the effects of these cosolutes on the dynamics of individual gene expression processes have not been systematically quantified. Here, we carry out kinetic mRNA and protein level measurements on libraries of genetic constructs using the common cosolutes PEG-8000, Ficoll-400, and magnesium glutamate. By combining these measurements with biophysical modeling, we show that cosolutes have differing effects on transcription initiation, translation initiation, and translation elongation rates with trade-offs between time-delays, expression tunability, and maximum expression productivity. We also confirm that biophysical models can predict translation initiation rates in TX-TL using *E. coli* lysate. We discuss how cosolute composition can be tuned to maximize performance across different cell-free applications, including biosensing, diagnostics, and biomanufacturing.

## Introduction

Cell-free expression systems combine in vitro transcription and translation (TX-TL) within a reconstituted cellular environment, enabling the expression of RNAs and proteins in an open biochemical system. Due to the ease of introducing novel components, and subsequent shortening of the design-build-test cycle, these systems have been harnessed across many synthetic biology applications. New genetic parts are rapidly characterized in organism-specific environments^1–5^. Enzymes can quickly be expressed or combined to prototype synthetic metabolic pathways with maximal productivity^6, 7^. Engineered state-switching RNAs and proteins are expressed, stored, and activated to detect viral nucleic acids, pollutants, and biomarkers of interest^8–11^. While the cell-free application space has grown considerably over the past few years, our understanding of how the cell-free environment affects genetic circuit and pathway function has not kept pace. This technical debt could limit future application development by reducing our ability to engineer more complex genetic systems inside cell-free environments.

Notably, the compositions of cell-free systems are chemically distinct from any *in vivo* environment. Overall, lysate-based cell-free systems are 20 to 30-fold more dilute than their corresponding cellular systems, though several salts, small molecules, and macromolecular crowding agents are added at much higher concentrations than typically found inside cells^12–15^. There are now several recipes for different cell-free systems, optimized for different purposes, making it difficult to interpret quantitative measurements and compare results across systems^12, 14^. Changes in solute composition have a poorly understood effect on the many steps in gene expression, particularly on the quantitative activities of promoters, ribosome binding sites, and other genetic parts^16–19^. Currently, it remains challenging to predict how these physio-chemical solute effects alter genetic part activities. For example, existing biophysical models of translation initiation rate do not take into account differences in solute composition^20, 21^. Consequently, the concentrations of these components are often empirically tuned to maximize a desired functionality, for example, *in vitro* protein expression titers or genetic circuit signal amplification^19, 22, 23^.

Here, as part of an effort to develop a more comprehensive mechanistic understanding of cell-free solvent effects, we systematically characterize how cell-free composition controls transcription, translation initiation, and translation elongation rates, as experimentally verified by dynamic mRNA level and protein level measurements. We developed a Markov model of translation that combines a statistical thermodynamic model of translation initiation (the RBS Calculator) with a thermodynamic model of solute-RNA interactions. We show that changes in cell-free composition can have differing effects on translation initiation versus elongation, leading to translation elongation becoming a rate-limiting step to protein expression. The developed model explains how changing the concentrations of commonly added solutes and crowding agents collectively control cell-free protein expression levels.

Finally, we suggest how TX-TL reaction compositions could be tuned for various cell-free applications, including genetic system prototyping, biomanufacturing, and sensing.

## Results & Discussion

### Kinetic Characterization of Genetic Systems in TX-TL with Varied Cosolute Compositions

We selected PEG-8000(PEG), Ficoll-400(Ficoll), and magnesium glutamate (Mg-glut) as three cosolutes commonly added to cell-free expression systems (**Figure 1A**). PEG is a polymer of ethylene glycol with an average molecular weight of 8 kDa and a hydrodynamic radius of about 2.6 nm^15^. Ficoll is a branched polysaccharide polymer with an average molecular weight of 400 kDa and a hydrodynamic radius of about 10 nm. Both PEG and Ficoll are crowding agents that reduce the total volume available to other macromolecules, greatly increasing effective concentrations inside cell-free expression reactions. Mg-glut is a commonly added salt that increases Mg^2+^ concentration, which has significant effects on nucleic acid interactions and mRNA folding.

**Figure 1.**
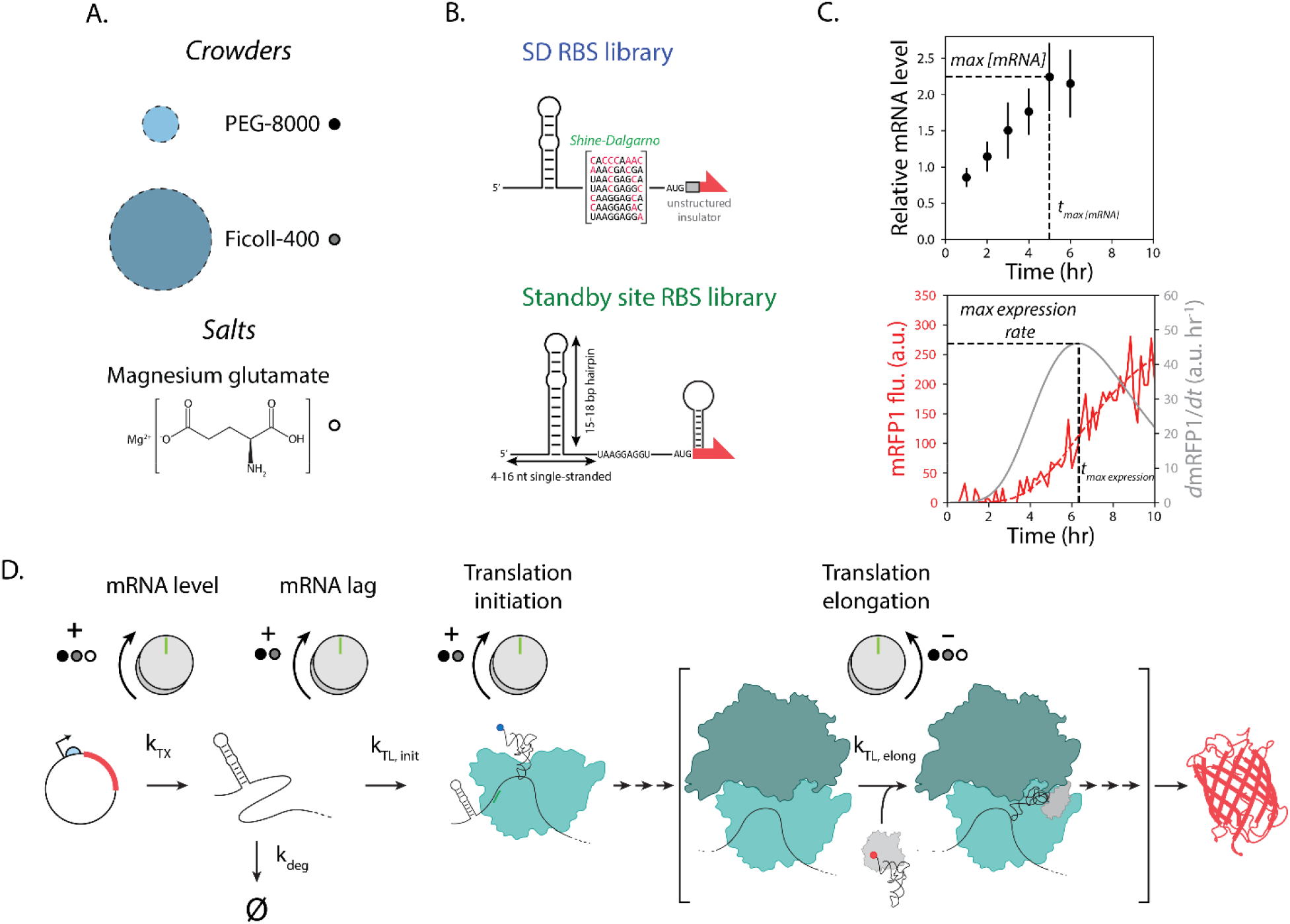
Measurements and models to characterize the effects of cosolutes on cell-free TX-TL assays.(A) Crowders and salts used in this study. (B) Sequence and structural differences for two ribosome binding site libraries. The SD RBS library varies translation initiation rates through changes to the Shine-Dalgarno sequence, while the Standby site RBS library varies translation initiation rates by altering the structural geometry of the upstream standby site. (C) The dynamics of mRNA and protein levels were measured using RT-qPCR and spectrophotometry, across several genetic constructs and cosolute compositions, during cell-free TX-TL assays. *mRFP1* expression dynamics were fitted to a 4-parameter logistic equation (red solid line). The first derivative of this equation (gray solid line) was used to determine the maximum expression rate. Error bars indicate 95% confidence intervals. (D) Biophysical modeling is used to distinguish the effects of cosolutes on mRNA dynamics, translation initiation rates, and translation elongation rates.

We then constructed a series of genetic systems expressing mRFP1 utilizing rationally designed ribosome binding sites (RBSs) with varied translation initiation rates. Two types of synthetic RBSs were designed, each varying a distinct interaction responsible for ribosome recruitment (**Figure 1B**). In the first set, the synthetic RBSs utilize different Shine-Dalgarno sequences with systematically varied hybridization energies to the 3’ end of the 16S rRNA, though they all contain an upstream insulating mRNA structure and an unstructured region at the beginning of the protein coding sequence (CDS). These SD RBS library variants were previously characterized using *in vivo E. coli* cultures, where they varied mRFP1 expression levels by 1649-fold with well-predicted translation initiation rates (RBS Calculator model v2.1; R^2^ = 0.99, p = 9×10^−7^)^24^. In the second set, the synthetic RBSs utilize different standby site sequences with varied structural geometries, followed by a canonical Shine-Dalgarno sequence (9 nucleotides long), an optimal spacer, and a small mRNA structure inside the beginning of the CDS. These Standby site RBS library variants were also previously characterized using *in vivo E. coli* cultures, varying mRFP1 expression by 40-fold with similarly well-predicted translation initiation rates (RBS Calculator model v2.1; R^2^ = 0.94, p = 4×10^−4^)^20^. All genetic systems utilized the J23100 promoter, which has a moderate transcription initiation rate.

The purpose of the Standby site RBS library variants is to vary how fast a 30S ribosomal subunit can initially bind to the mRNAs’ standby sites. Once a 30S ribosomal subunit is bound, these RBSs all have strong 16S rRNA binding sites, facilitating a rapid transition to forming a 30S pre-initiation complex (PIC) and initiating translation. In contrast, the SD RBS library variants all have a highly accessible upstream standby site, but the differences in their 16S rRNA binding sites lead to different transition rates in 30S PIC formation. Overall, the Standby site RBS library varies the rate of the first key step of translation initiation (a 2^nd^ order binding event) whereas the SD RBS library varies the rate of the second key step in translation initiation (a 1^st^ order sliding event).

For each of these genetic systems, we then carried out kinetic mRNA level and mRFP1 fluorescence level measurements during 10-hour cell-free expression assays, adding 2 nM circular plasmid template, and using baseline or systematically varied cosolute compositions (**Methods**) (**Figure 1C**). For each cosolute composition, mRNA level measurements were performed (N = 3 biological replicates) using RT-qPCR with a temporal resolution of 1 hour. mRFP1 fluorescence level measurements were performed (N = 6 biological replicates) using spectrophotometry with a temporal resolution of 10 minutes. Notably, during our RT-qPCR measurements, we found that endogenous 16S rRNA degrades over time. Therefore, as our internal control, we instead added a synthetic spike-in RNA at fixed concentration to each reaction immediately before RNA extraction (**Methods**). From these measurements, we determined the dynamics of the mRNA and protein levels, including the time to reach maximum mRNA levels, the first derivative (slope) of the mRFP1 fluorescence levels, and the apparent translation rates of the mRNAs. For the purpose of slope calculations, we found that mRFP1 fluorescence levels fit well to a generalized logistic growth curve, which accounts for background autofluorescence and time delays (**Methods**). From these measurements, we determined how the concentrations of PEG, Ficoll, and Mg-glut differentially controlled the genetic system variants’ expression dynamics, including delays in mRNA synthesis, maximum mRNA levels, translation initiation rates, and translation elongation rates (**Figure 1D**).

### Cosolute Composition Differentially Controls Dynamics of mRNA and Protein Levels

We first measured the mRNA level dynamics of two selected genetic system variants in TX-TL reactions with modified compositions of 4% w/v PEG, 4% w/v Ficoll, or 16.67 mM Mg-glut, as compared to the baseline composition. The baseline solution contains 8.67 mM Mg-glut, as TX-TL reactions containing less Mg-glut were unable to support robust expression from the weaker RBSs, but does not contain any PEG or Ficoll. The selected genetic system variants included a Standby Site RBS library variant and a SD RBS library variant with similar predicted translation initiation rates. Notably, we did not find any appreciable difference in mRNA dynamics across these genetic system variants. However, we found that changing the cosolute composition had distinct effects on transcription delays and mRNA maximum levels (**Figure 2A**). Overall, PEG had the highest impact on altering the mRNA maximum level, increasing it by about 6.8-fold, followed by Mg-glut (2.5-fold) and Ficoll (1.9-fold). PEG also had the highest impact on variability in maximum mRNA levels across replicates; its average coefficient of variation across all time points was 0.53, which is about 2-fold higher than the baseline composition and other tested cosolutes. Interestingly, the addition of PEG or Ficoll increased the time needed to reach maximum mRNA levels, due to an apparent delay in mRNA synthesis. The time delay was about 2.5 hours for PEG and 1.3 hours for Ficoll as compared to the baseline composition. In contrast, the addition of Mg-glut did not cause any appreciable difference in time delay.

**Figure 2.**
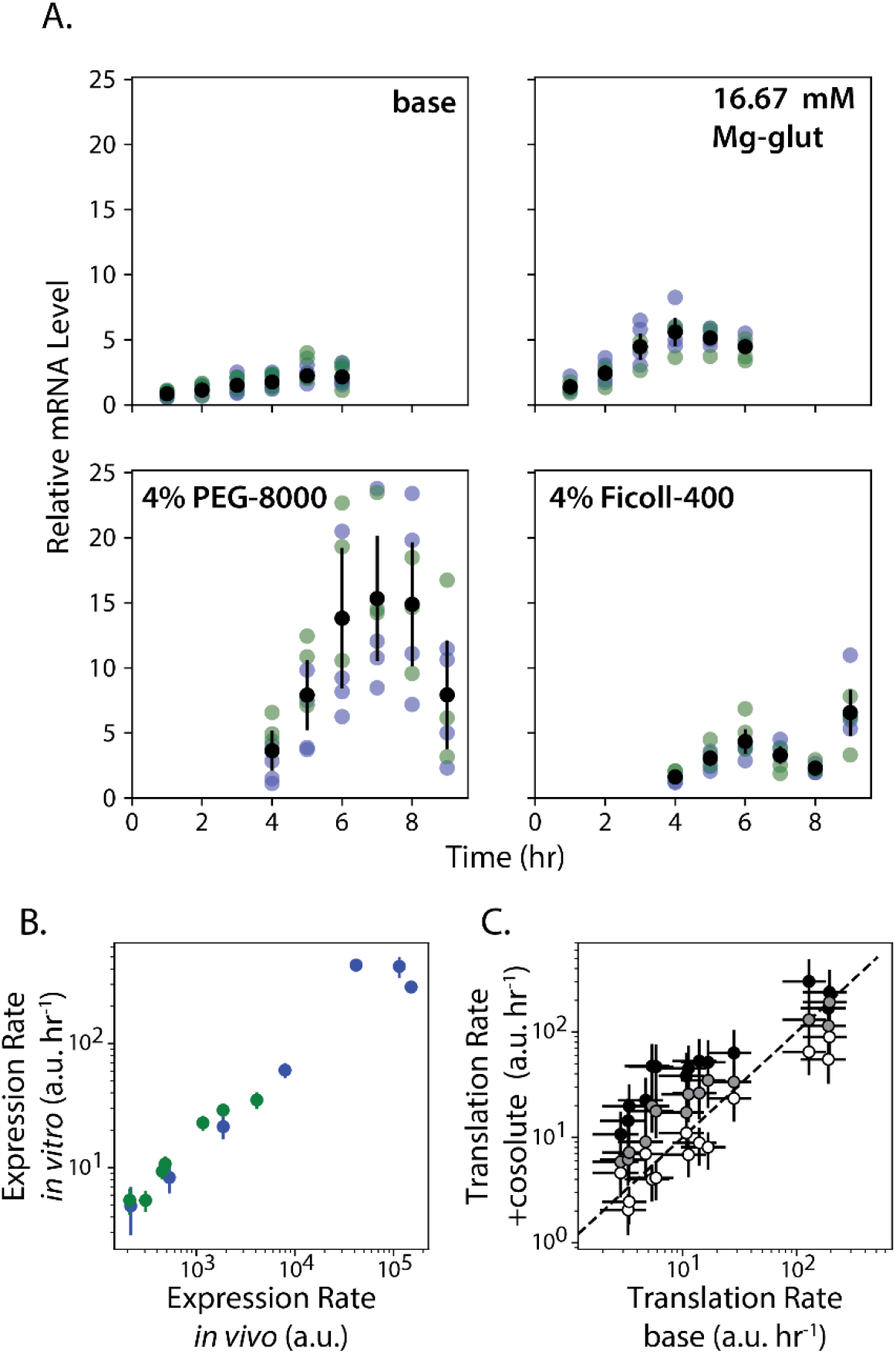
Cosolute effects on mRNA levels and translation rates. (A) Relative mRNA levels were measured when using 4% PEG-8000, 4% Ficoll-400, or 16.67 mM Mg-glut, as compared to a baseline composition. Data points are replicate mRNA level measurements from a SD RBS library variant (blue), a Standby Site RBS library variant (green), or the average across all replicates from both RBS library variants (black circles). Error bars represent the 95% confidence interval of at least 7 biological replicates. (B) Measured *in vivo* expression levels were compared to cell-free expression levels across SD RBS library variants (blue) and Standby site RBS library variants (green). (C) Translation rates for all RBS library variants at various cosolute compositions were compared to the translation rates at the baseline composition. Data points include 4% PEG-8000 (black circles), 4% Ficoll-400 (gray circles), and 16.67 mM Mg-glut (white circles). The dotted line indicates no differences. For (B) and (C), data points and error bars represent the mean and 95% confidence interval of at least 6 biological replicates.

We then measured the maximum synthesis rates of *mRFP1* mRNA and protein for all genetic system variants – 7 Standby Site RBS library variants and 7 SD RBS library variants – during TX-TL reactions, using modified compositions of either 4% w/v PEG, 4% w/v Ficoll, or 16.67 mM Mg-glut, as compared to the baseline composition. We also measured these genetic system variants’ *in vivo* mRFP1 expression levels in *E. coli* DH10B during steady-state cultures maintained in the exponential growth phase (**Methods**). Notably, at the baseline TX-TL composition, the genetic system variants’ protein synthesis rates (expression rates) were highly correlated to their *in vivo* expression levels (R^2^ = 0.96, p = 5×10^−10^, **Figure 2B**), though the dynamic ranges were starkly different. Changing the RBS sequence varied *in vivo* expression levels by 717-fold, while the same RBS sequences varied in vitro expression levels by only 67.9-fold.

We then examined how cosolute composition affected the genetic system variants’ apparent translation rates. To calculate the apparent translation rates, we divided the measured protein synthesis rates by the measured mRNA levels. Surprisingly, we found that adding PEG or Ficoll greatly distorted the tunability of apparent translation rates as compared to the baseline composition or *in vivo* measurements. RBS library variants with the lowest measured translation rates (at the baseline composition) had the highest increases in protein synthesis rates (8.8-fold when adding 4% w/v PEG or 3.7-fold when adding 4% w/v Ficoll) (**Figure 2C)**. However, this distortion was diminished when using RBS library variants with high measured translation rates. In contrast, adding additional Mg-glut had no appreciable effect on the RBSs’ translation rates, which remained correlated with their *in vivo* expression levels. Overall, depending on the cosolute added to TX-TL, there are extremely large changes in maximum mRNA levels, delays in mRNA synthesis, apparent translation rates, and expression tunability.

### Cosolute Composition Controls the Magnitude and Timing of Protein Expression Levels

Our next objective was to systematically vary cosolute composition and quantify their effects on the magnitude, timing, and tunability of protein expression levels across all 14 genetic system variants with varied RBS sequences. We first found that increasing PEG from 0 to 4% w/v greatly increased maximum protein expression levels by an average of 27.6-fold (**Figure 3A**, top). Similar to our previous measurements of the RBS variants’ apparent translation rates, the cosolute effect was most pronounced on the RBS library variants with the lowest translation rates. As a result, the addition of PEG also greatly reduced the dynamic range of expression tunability by 3.1-fold. This effect was significant even at 1% w/v PEG and was further enhanced at higher PEG concentrations. As before, the addition of Ficoll yielded a similar effect with a smaller magnitude; 4% w/v Ficoll increased maximum protein expression levels by an average of 3.6-fold and reduced the dynamic range of tunability by 2.1-fold (**Figure 3A**, middle). Interestingly, Mg-glut only increased maximum protein expression levels by 1.9-fold, but decreased expression tunability by a larger amount (2.8-fold) (**Figure 3A**, bottom). The timing of expression was also significantly affected by cosolute composition. Systematic increases in PEG concentration increased the time needed to reach maximum protein expression by 2.5 hours (**Figure 3B**, top). Similarly, increasing Ficoll concentration resulted in a delay of 1.3 hours to reach maximum protein expression levels (**Figure 3B**, middle). In contrast, changing the Mg-glut concentration had little appreciable effect on overall expression timing (**Figure 3B**, bottom).

**Figure 3.**
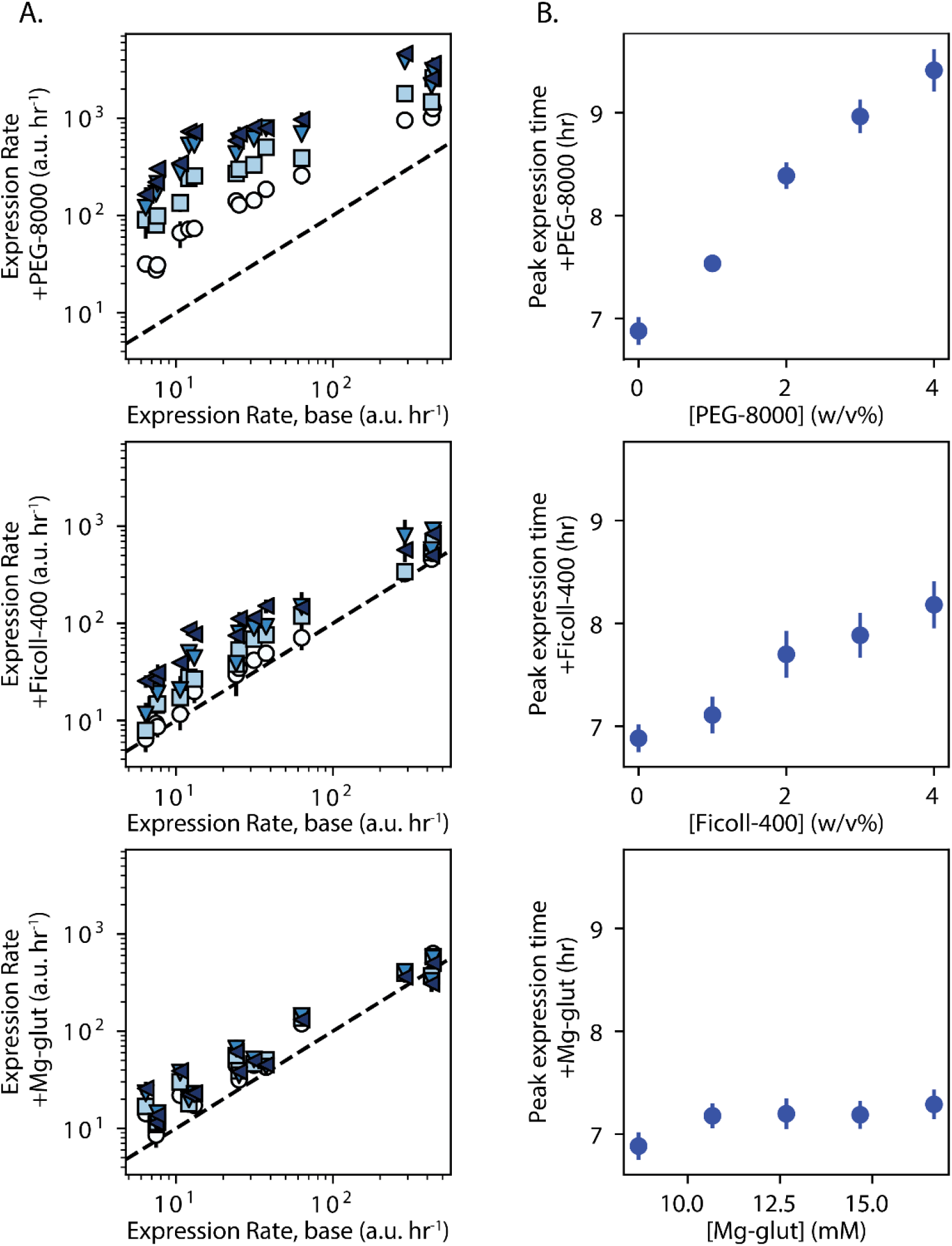
The effects of tuning cosolute composition on the maximum expression rate and peak expression time in cell-free assays. (A) The maximum expression rate was determined for all RBS library variants using either 1%, 2%, or 3%, or 4% PEG-8000 (top); 1%, 2%, 3%, or 4% Ficoll-400 (middle); or 10.67, 12.67, 14.67, or 16.67 mM Mg-glut (bottom). Data points of increasing composition are white circles, light blue squares, blue down-triangles, or dark blue up-triangles. Data points and error bars represent the mean and 95% confidence interval of at least 6 biological replicates. (B) The average peak expression time for all tested RBSs was determined when varying cosolute compositions. Data points and error bars represent the mean and 95% confidence interval for 14 genetic systems with 6 biological replicates each.

### Biophysical Modeling Explains Changes in Translation Initiation and Elongation Rates

We next investigated how biophysical modeling can explain both the sequence-dependent and cosolute-dependent effects on translation initiation and elongation rates. As a baseline, we found that the mRFP1 expression levels from the 14 genetic system variants, as measured *in vivo* within *E. coli* DH10B cells, were highly proportional to the RBS Calculator v2.1 model’s predicted translation initiation rates, suggesting that translation initiation was a key rate-limiting to protein production (**Figure 4A**, R^2^ = 0.813, p = 1×10^−5^, N = 14). However, when expressing the same genetic systems in TX-TL, we found that the level of proportionality was reduced when adjusting the composition to either baseline, 4% w/v PEG, 4% w/v Ficoll, or 16.67 mM Mg-glut (overall R^2^ = 0.63, p = 2×10^−13^, N = 64 conditions, **Figure 4E**). Overall, we found that the cosolute composition had a large impact on the proportionality constant relating model-predicted translation initiation rates to measured expression levels. We also observed a plateau effect whereby higher translation initiation rates did not yield appreciably higher mRFP1 expression levels (**Figure 4E**). Together with our prior measurements (**Figure 2**), these observations suggested that the composition of the TX-TL reaction has distinct effects on the magnitudes of both the translation initiation and translation elongation steps, potentially making translation elongation a rate-limiting step during protein expression.

**Figure 4.**
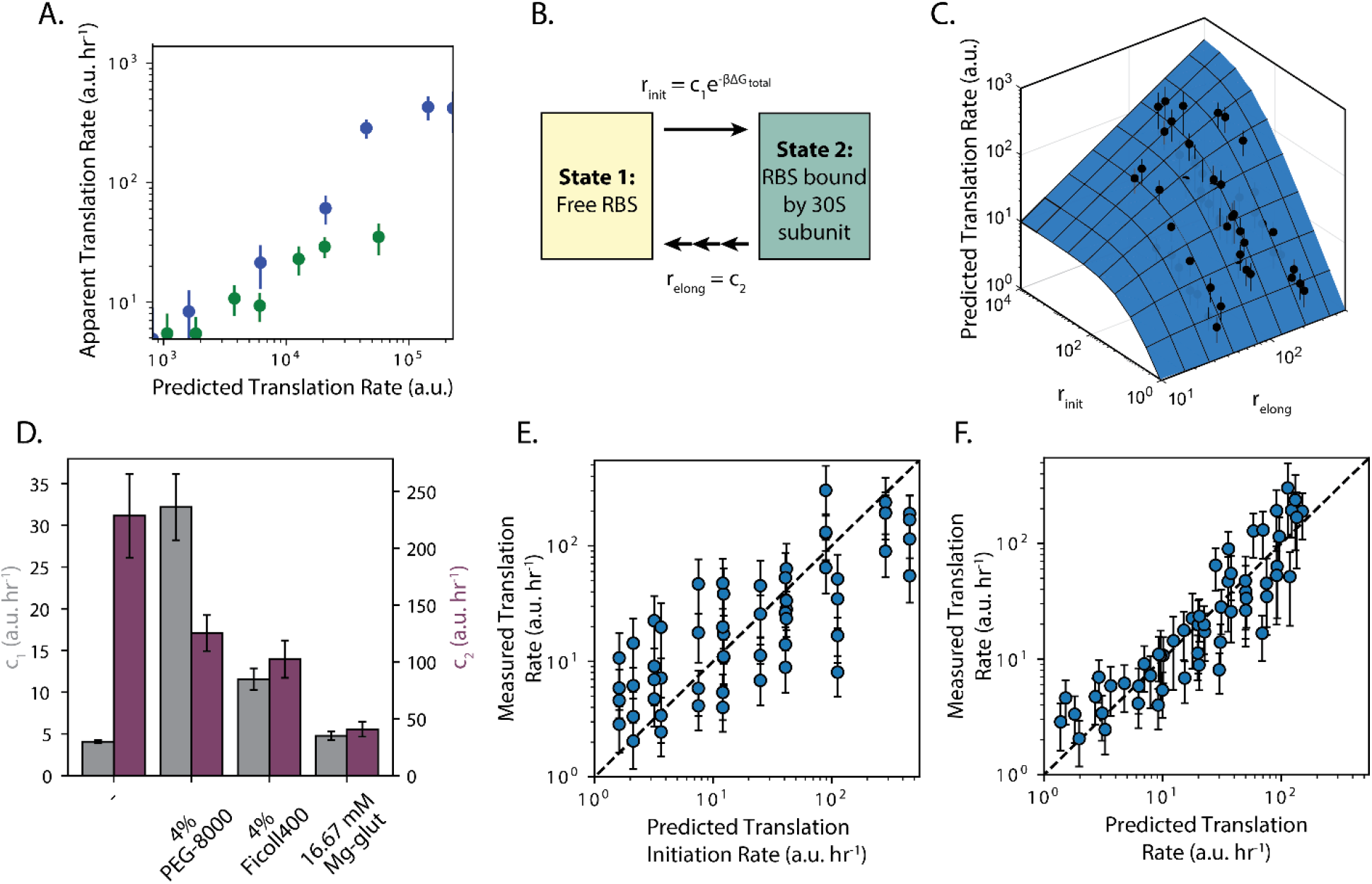
Biophysical modeling quantifies effects of cosolutes on gene expression processes. (A) Predicted translation initiation rates were compared to measured *in vivo* translation rates for SD RBS library variants (blue) and Standby Site RBS library variants (green). (B) A simplified two-state model is used to distinguish cosolute effects on translation initiation and translation elongation steps. (C) The parameterized model shows how cosolute-mediated changes in translation initiation and translation elongation control protein expression levels. (D) Measurements were used to identify parameter values c_1_ (gray bars) and c_2_ (purple bars), which quantify cosolute effects on translation initiation and translation elongation. Error bars are 95% confidence intervals for the fitted parameters. (E) Measured translation rates are insufficiently predicted by translation initiation rate alone *in vitro*. (F) Accounting for cosolute effects on both translation initiation and translation elongation rates increases predictive accuracy of model. Circles and error bars for (A), (C), (E), and (F) represent the mean and 95% confidence interval of at least 6 measurements.

We therefore augmented the RBS Calculator model to explicitly include the apparent translation elongation rate of the protein’s coding sequence. As the starting point, the RBS Calculator calculates the ribosome’s binding free energy (ΔG_total_) using a 5-term free energy model^20, 21, 25, 26^ and then predicts a protein coding sequence’s translation initiation rate (r_init_) according to:

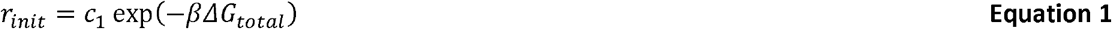

where the 30S ribosomal subunit’s binding free energy (ΔG_total_) depends only on the mRNA’s sequence and c_1_ is a proportionality constant that accounts for extrinsic differences here influenced by the cosolute composition. After the 30S ribosomal subunit binds to the mRNA, it recruits the 50S ribosomal subunit, forms the 70S initiation complex, and initiates translation. Translation continues with a highly processive cycle with elongation rates that depend on codon identities, charged tRNA availabilities, and the cosolute composition. As soon as an elongating ribosome clears the ribosome binding site, a new 30S ribosomal subunit may bind to initiate a new cycle of translation, leading to polysome multi-ribosome dynamics.

Here, our objective is to determine how the cosolute composition affects the translation elongation rate, averaged over the codons in the mRFP1 protein coding sequence. We designated this averaged translation elongation rate as r_elong_ and formulated the simplest possible two-state model (**Figure 4B**) that accounts for how r_init_ and r_elong_ together control the translation rate (r_TL_), according to the equation:

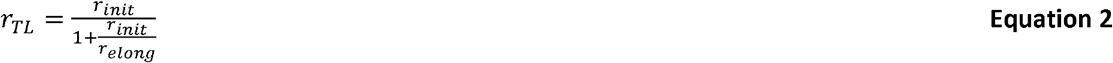

where r_elong_ is c_2_, a single cosolute-dependent coefficient that quantifies how the cosolute composition affects the average translation elongation rate of the mRFP1 coding sequence.

We then carried out a model identification procedure to determine the coefficients c_1_ and c_2_ for each cosolute condition (**Methods**), yielding a mean and confidence interval for each coefficient value. The parameterized two-state model (**Equation 2**) clearly shows how cosolute composition can negatively affect translation elongation rates, resulting in lower translation rates than otherwise predicted by the RBS Calculator model (**Figure 4C**). Overall, by including translation elongation, the two-state model is now able to accurately predict mRFP1 expression levels across all 14 genetic systems and 4 cosolute compositions (R^2^ = 0.81, p = 4×10^−21^, N = 64, **Figure 4F**).

More importantly, the fitted model shows us how each cosolute affected each translation step, providing an explanation for the observed phenomenon. For example, 4% w/v PEG and 4% w/v Ficoll increased the apparent translation initiation rates by 7.94-fold and 2.85-fold, respectively (**Figure 4D**). In contrast, 16.67 mM Mg-glut did not appreciably change the initiation rate (1.18-fold increase), consistent with our prior measurements (**Figure 3**). Surprisingly, all of the cosolutes had a negative impact on translation elongation rates. 16.67 mM Mg-glut had the largest effect; it lowered the apparent translation elongation rate by 5.6-fold whereas 4% w/v PEG and 4% w/v Ficoll lowered it by 1.82-fold and 2.23-fold, respectively. As a result, the model shows why adding a cosolute decreases the overall expression tunability when utilizing different RBS sequences. For 4% w/v PEG, the model shows that the increase in initiation rate and decrease in elongation rate yields the observed plateau effect whereby RBSs that bind better to ribosomes (strong RBSs) do not yield appreciably more protein than weak RBSs. For 4% w/v Ficoll, a similar plateau effect is predicted, though with lower overall amounts of expressed protein. For 16.67 mM Mg-glut, the model shows that expression tunability is limited by primarily making translation elongation a rate-limiting step in protein production.

### Physical Modeling Connects Cosolute Intrinsic Characteristics to Extrinsic Effects on Translation

To better understand why PEG, Ficoll, and Mg-glut can have such distinct effects on gene expression, we next applied physical modeling to calculate how cosolute composition affects the kinetic and thermodynamic properties of these non-ideal liquids (**Figure 5**). We first investigated how PEG and Ficoll composition affects the rate of diffusion inside TX-TL reactions. The rate of diffusion affects all binding interactions inside TX-TL, but will particularly affect the charging and loading of tRNAs during translation elongation, which are the slowest and most diffusion-limited steps in protein expression^27, 28^.

**Figure 5.**
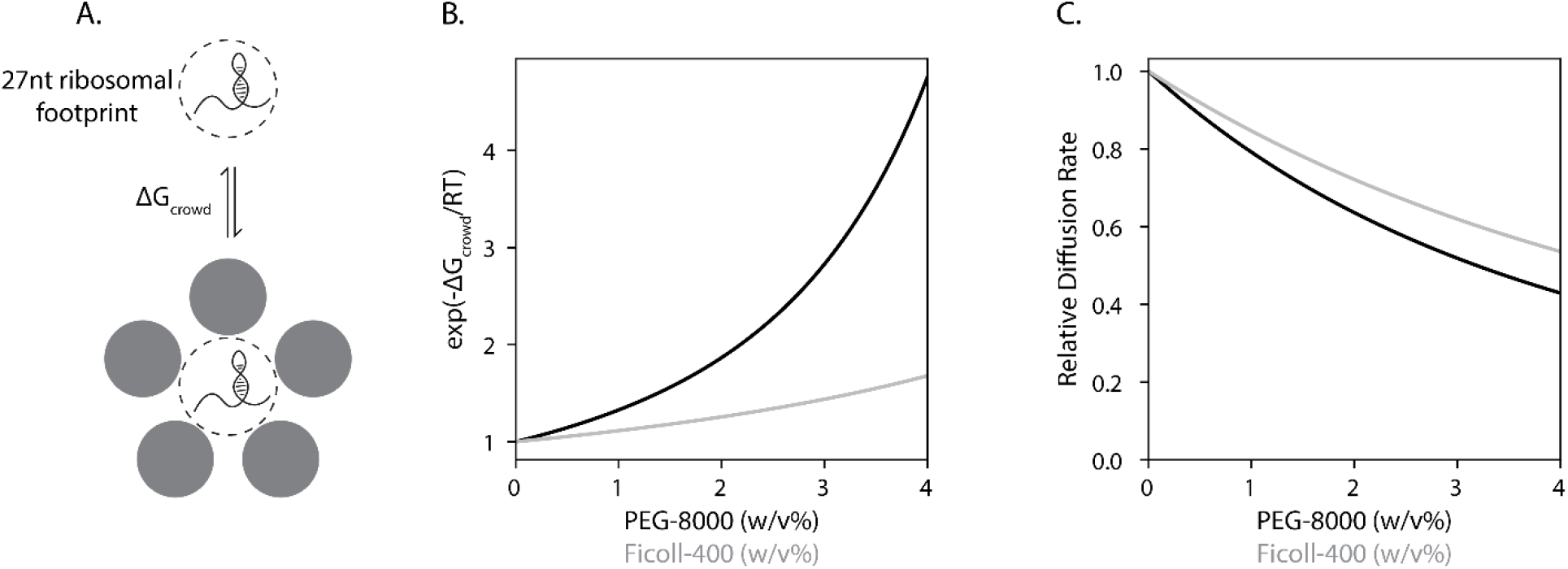
Crowding affects both translation initiation and molecular diffusion. (A) PEG and Ficoll bind to the 27-nt ribosomal footprint region of an mRNA with free energy ΔG_crowd_, which increases a ribosome’s ability to bind and initiate translation. (B) Scaled Particle Theory is used to calculate the enhancement of translation rates by varying PEG-8000 (black) and Ficoll-400 (gray) concentrations. (C) The Stokes-Einstein-Huggins equation is used to calculate the decrease in relative diffusion rates when varying PEG-8000 (black) and Ficoll-400 (gray) concentrations.

To do this, we combined the Stokes-Einstein equation with the Huggins equation to calculate how a cosolute’s intrinsic viscosity and its concentration control the solution’s diffusion coefficient, yielding:

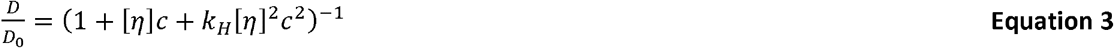

where *D* is the diffusion coefficient of the TX-TL solution with cosolute present at concentration c, D_o_ is the diffusion coefficient in the absence of cosolute, [*η*] is the intrinsic viscosity of the cosolute (in a dilute solution), and k_H_ is the Huggins coefficient of the cosolute. The intrinsic viscosities of PEG and Ficoll are 17 mL/g and 24 mL/g, respectively^29^. Typical Huggins coefficients vary from 0.3 to 0.5; here, in the absence of direct measurements, we assume that PEG and Ficoll both have a Huggins coefficient of 0.4. Using **Equation 3**, we calculated that 4% w/v PEG and 4% w/v Ficoll reduce the TX-TL diffusion coefficient by 2.2-fold and 1.8-fold, respectively (**Figure 5C**). This reduction in the diffusion coefficient is quantitatively similar to the observed reduction in translation elongation rates for both PEG and Ficoll (**Figure 4D**).

Next, we considered how the divalent salt Mg-glut affects interactions during TX-TL expression. Notably, it has been determined that excess amounts of free Mg^2+^ will inhibit tRNA translocation through the 70S ribosome during translation elongation^30^. The amount of free Mg^2+^ is greatly determined by the concentration of other metabolites in TX-TL that act as chelators, such as glutamate (K_d_ = 15.1 mM)^31^ and phosphoenolpyruvate (K_d_ = 11 mM)^32^. Taking into account the concentrations of these chelators, we calculate that the concentration of free Mg^2+^ is 0.79 mM in the baseline TX-TL composition (no added Mg-glut). The free Mg^2+^ then increases to 1.45 mM when adding additional Mg-glut to 16.67 mM. Doubling the free Mg^2+^ concentration results in at least a 2-fold reduction in ribosome translocation rate^30^, suggesting a causal mechanism for the observed inhibition of translation elongation at high Mg-glut concentrations (**Figure 4D**).

Finally, we considered how PEG and Ficoll affect the thermodynamics of ribosome-mRNA interactions during translation initiation. As crowding agents, PEG and Ficoll interact with other chemical components in solution, reducing the amount of free volume available to them, and promoting the formation of more compact states that take up less space. In this way, the magnitude of the crowding effect depends on the size and shape of the interacting components, particularly larger macromolecules (e.g. mRNAs). Qualitatively, the addition of a crowding agent favors the formation of the 30S:mRNA complex as the bound state takes up less space in solution as compared to a free 30S subunit and a free mRNA. Here, we leveraged Scaled Particle Theory (SPT)^33^ to quantitatively calculate the thermodynamic free energies between crowding agents and solution components to determine the magnitude of this crowding effect. To do this, we first assume that each particle is a hard body sphere and then leverage prior measurements to determine their sizes. The cosolutes each have a defined radius (PEG R_c_ = 2.6 nm, Ficoll R_c_ = 10 nm), and vary in volume fraction (ϕ_c_, unitless) in a composition^34^. As a particle, the 30S ribosomal subunit has a radius of about R_m_ = 11 nm^35, 36^. We then consider only the portion of the mRNA that binds to the 30S ribosomal subunit during translation initiation – a 27-nucleotide region called the ribosome footprint – and its defined radius (R_m_ = 1.75 nm)^37^. We then use SPT to calculate the positive free energy (ΔG_SPT_) when crowding agent binds to either the 30S ribosomal subunit, the 27-nt mRNA region, or the bound 30S:mRNA complex, according to the equation:

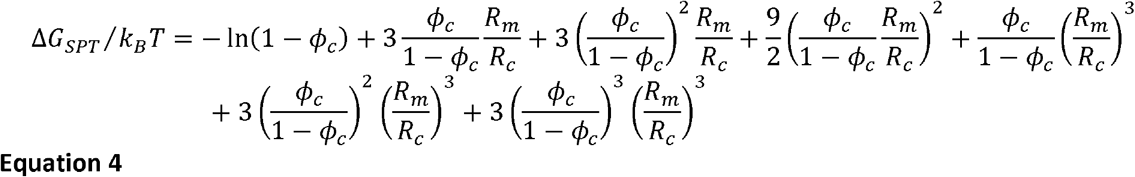

where k_B_T is the Boltzmann constant multiplied by the system temperature. ΔG_SPT_ is positive as it requires an input of energy to add a 30S or mRNA particle into a crowded solution with limited free space. We then calculate the difference in free energy when considering a 30S:mRNA particle added to a crowded solution versus a free 30S subunit and a free mRNA added to a crowded solution, according to Equation 5:

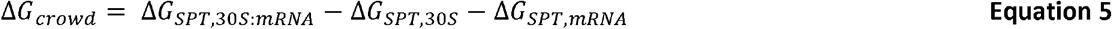

The quantity ΔG_crowd_ is the negative free energy that quantifies how the addition of crowding agent promotes the formation of the more compact 30S:mRNA complex. Because the ribosome is so much larger than the mRNA, here we assume that the 30S:mRNA particle has the same radius as the 30S particle. Therefore, ΔG_crowd_ simplifies to −ΔG_SPT,mRNA_. In Figure 5B, we show how ΔG_crowd_ becomes more negative as more crowding agent is added, leading to higher translation initiation rates. We calculate that 4% PEG and 4% Ficoll increase a mRNA’s translation initiation rate by 4.8-fold and 1.7-fold, respectively. These calculations agree with our empirical measurements shown in **Figure 4D**.

## Discussion

Cell-free expression systems are now commonly used for biomanufacturing, diagnostic assays, and genetic part characterization^1, 2, 8, 38–44^. However, the amounts of cosolute added to prepared cell-free assays can vary considerably across protocols, without a clear understanding of how cosolute composition will affect the performance metrics for each of these applications^14, 45^. For example, when using cell-free expression to manufacture proteins, it is highly desirable to adjust cosolute concentration to maximize the amount of protein produced, though it less important to activate protein production as rapidly as possible^46^. In contrast, when using cell-free expression as a diagnostic assay, it is more important to rapidly activate production of the observable output^42^. Expression tunability is another important performance metric when multiple proteins need to be expressed at different rates.

Here, we investigated how the concentrations of three cosolutes (PEG, Ficoll, and magnesium glutamate) affected the dynamics of transcription, translation initiation, and translation elongation across 14 genetic systems with varied genetic parts. Overall, we found that all cosolutes increased transcription rates, with PEG having the highest impact. All cosolutes decreased translation elongation rates, with Mg-glut having the highest impact. However, only PEG and Ficoll increased translation initiation rates with PEG having the highest impact. Altogether, the addition of cosolutes increased protein synthesis rates, though the magnitude of improvement depended on which genetic parts were used. When using weak ribosome binding sites, the addition of PEG or Ficoll increased the apparent translation initiation rate, resulting in higher protein expression. However, when using stronger ribosome binding sites, the addition of any cosolute lowered the translation elongation rate, causing it to become the rate-limiting step during protein production and creating a plateau in the protein synthesis rate. We then applied theory from physical chemistry to explain how the cosolutes’ intrinsic differences in viscosity, size, and charge could be responsible for these effects through alteration of the cell-free assay’s solvent properties.

From these results, we observe distinct trade-offs between timing, expression tunability, and maximum protein production that affect how cosolute composition directly impacts an application’s performance metrics (**Figure 6**). Adding PEG or Ficoll will lead to much higher protein production rates, though at the cost of introducing a substantial time delay and limiting the ability to tune protein expression levels by varying translation rates. In contrast, Mg-glut has a much smaller effect on all these performance criteria. With these trade-offs in mind, it becomes possible to rationally tune cosolute composition towards maximizing performance metrics for a particular application. In a biomanufacturing application where only a single protein needs to be expressed, adding a large amount of PEG will increase the overall protein production rate and titer. However, when multiple proteins need to be expressed at different levels, the PEG and Ficoll concentrations can be tuned to achieve the minimum level of expression tunability, while maximizing the overall protein production rates. In contrast, for a diagnostic application where the time delay becomes more important, the absence of any additional cosolute may instead be the optimal choice.

**Figure 6.**
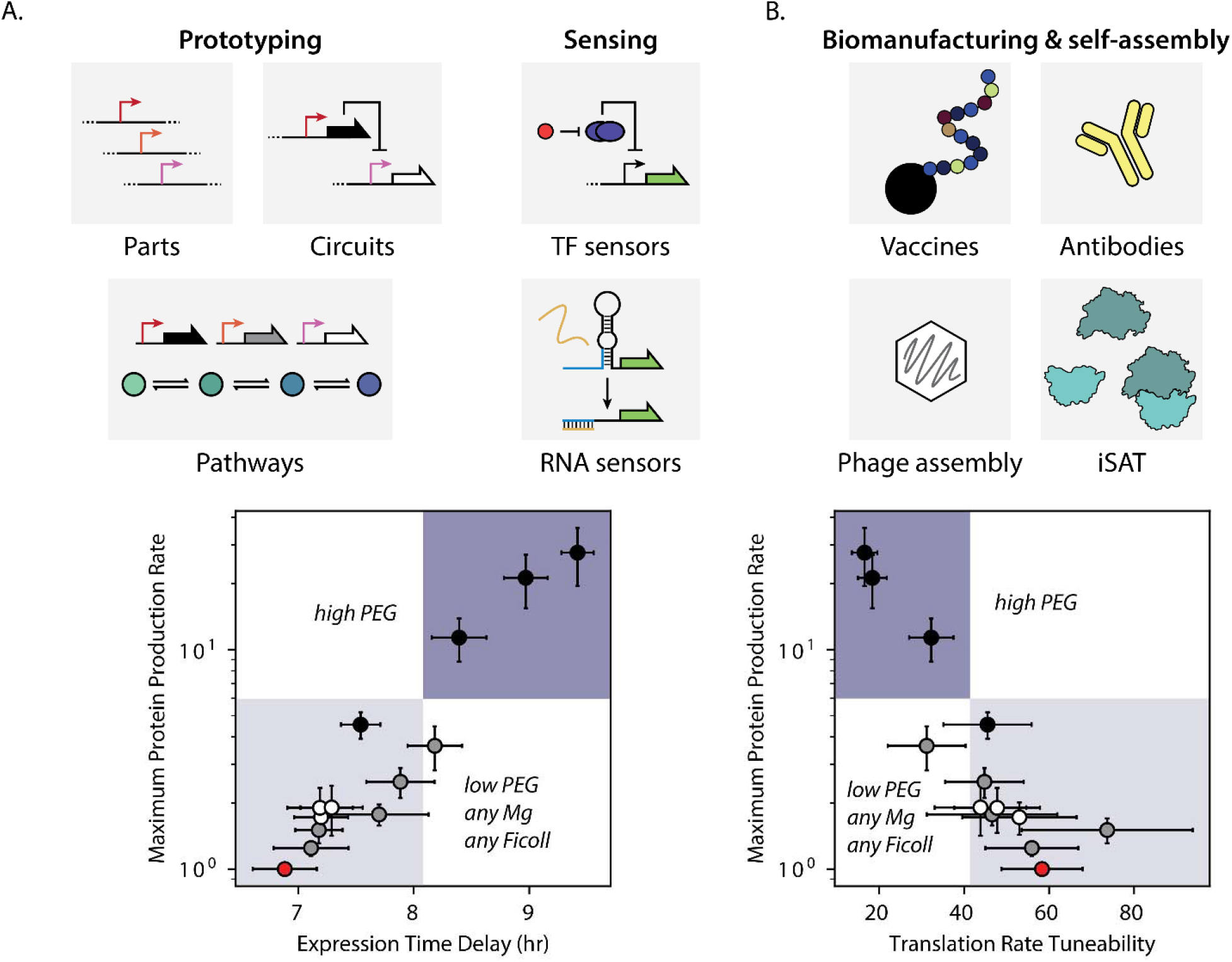
Trade-offs between cell-free expression timing, tunability, and magnitude. (A) Several cell-free applications require rapid turn-on times, high tunability, and high dynamic range in expression levels. When tuning cell-free composition, there is a trade-off between a rapid turn-on time and expression tunability. (B) Several cell-free applications require high production rates of protein. Here, cell-free composition can be tuned to maximize protein expression, though sacrificing expression tunability. Circles in (A) and (B) are base composition (red) or increasing concentrations of PEG (black), Ficoll (gray), or Mg-glut (white). Circles and error bars represent the mean and 95% confidence interval at least 6 biological replicates.

Finally, our results show that it can be highly misleading to utilize cell-free systems to characterize genetic parts for later re-use in *in vivo* systems. The cosolute composition of all cell-free systems are distinct from the *in vivo* environment, and these differences play an important role in controlling transcription and translation rates. First, differences in genetic part activities (e.g. the translation rates of ribosome binding sites) are often compressed inside cell-free expression systems, leading to lower changes in protein expression levels than otherwise expected. Second, because cosolutes decrease a mRNA’s translation elongation rate, the role of synonymous codon usage in protein coding sequences could play an even greater role in cell-free expression systems, particularly when using strong ribosome binding sites and/or high PEG/Ficoll concentrations. While we can apply theory from physical chemistry to calculate and predict how cosolute composition controls genetic part activities in cell-free systems, we should never assume that *in vivo* and cell-free systems will yield quantitatively equivalent results.

## Methods

### Crude cell lysate preparation

Crude cell lysate was prepared according to a previously published protocol, with the following modifications^12^. 20 L of Escherichia coli BL21 with the Rosetta2 plasmid encoding rare tRNAs was cultured in a Micros 30-liter fermentor (New Brunswick) in 2XYT+P medium until the cells reached an OD600 of 1.5-2.0. The cell pellet was then collected in a T-1-P Laboratory continuous flow centrifuge (Sharples), and resuspended in 1 mL S30A buffer per gram of cell pellet. The resuspended cells were run through a M110-EH-30 microfluidizer (Microfluidics Corp.) at 20,000 PSI twice to ensure complete lysis. The lysate was clarified by centrifugation at 12,000xg for 30 minutes at 4C. The clarified lysate was then incubated for 80 minutes at 37 minutes while undergoing orbital shaking to perform the runoff reaction. After incubation, the lysate was centrifuged again for at 12,000xg for 30 minutes at 4C.

Following lysis, clarification, and the runoff reaction, the lysate was diafiltered with a Pellicon Biomax 10 kDa MWCO 0.005 m^2^ ultrafiltration module. Six retentate volumes of buffer S30B were run against the lysate at 4C. After diafiltration, the retentate was centrifuged for 30 minutes at 12,000xg at 4C. The protein concentration of the retentate was quantified using a Bradford BSA Protein Assay Kit assay (Bio-Rad). The retentate was aliquoted and flash-frozen in liquid nitrogen, and stored at -80C.

### Cell-free expression reactions

Cell-free expression reactions were assembled on ice according to previously published protcols, with the following modifications^12, 45^. Amino acid and energy solutions were prepared separately, and combined with crude cell extract to reach the following final concentrations: 7.4 mg/mL protein (1/3^rd^ total reaction volume), 1.5 mM each amino acid (except for leucine at 1.25 mM), 50 mM HEPES, 1.5 mM ATP and GTP, 0.9 mM CTP and UTP, 0.2 mg/ml tRNA, 0.26 mM CoA, 0.33 mM NAD, 0.75 mM cAMP, 0.068 mM folinic acid, 1 mM putrescine, and 30 mM PEP. Unless otherwise indicated, 4 mM additional magnesium glutamate *(8.67 mM total), 80 mM additional potassium glutamate (100 mM total), and no crowding agents were added to each reaction. Plasmid DNA containing an RBS variant controlling the expression of an mRFP1 reporter was either miniprepped and ethanol precipitated, or midiprepped and isopropanol precipitated, and added to the reaction to a final concentration of 2 nM. 5 uL reactions were incubated at 29°C for 12 hours in a 96-well polypropylene conical bottom plate sealed with a plate storage mat (Corning) in a TECAN Spark microplate reader. mRFP1 fluorescence was measured every 10 minutes, using 584nm/60nm ex/em with a 5 nm bandwidth.

### RT-qPCR

5 uL TXTL reactions, assembled as above, were incubated at 29C in 96-well Costar conical-bottom plates in a TECAN Spark.After incubating for the given amount of time, reactions in microcentrifuge tubes were flash-frozen in liquid nitrogen, while reactions in 96-well plates were directly processed. To each reaction, 500 pM iclR normalization control RNA was. Total RNA, including the normalization control, was extracted using a Norgen Total RNA Extraction kit, after which any remaining plasmid DNA was removed via digestion with TurboDNase (Invitrogen). RNA integrity was verified via agarose gel electrophoresis. RNA was then diluted to 100 ng/uL, and 25 ng/uL yeast tRNA was added. First-strand cDNA synthesis was performed using a High-Capacity cDNA Reverse Transcription Kit (Applied Biosystems). qPCR reactions were assembled using PowerUp SYBR Green Master Mix (Applied Biosystems) and primer sets for mRFP1 and iclR, and run in a StepOnePlus Real-Time PCR System (Applied Biosystems). Melting curve analysis was performed to confirm product homogeneity. RNA levels were calculated using a variant on the ddCt method, accounting for differences in probe efficiency.

### Fluorescence time-course data analysis

To determine the time-course rates of gene expression, we fit the fluorescence production timecourse for each TXTL reaction to a generalized logistic growth model of the form^47^:

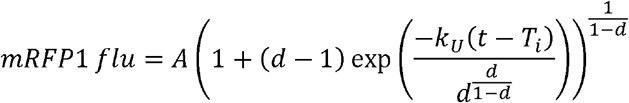

Where A is the upper asymptote, T_i_ is the time at inflection, k_U_ is the relative maximum growth rate, and d is a parameter controlling the inflection value. The generalized logistic curve allows for a variable amount of asymmetry about the inflection point, which better describes the behavior of fluorescent protein production over time in the cell-free expression system used here. Timecourses were corrected for background fluorescence by subtracting out the average fluorescence from 30-80 minutes. Fitting was performed using scipy’s curve_fit module. The rate of fluorescence production was calculated as the gradient of the fit fluorescence timecourse.

## Author Contributions

G.E.V. and H.M.S designed the study, analyzed results, developed models, and wrote the manuscript. G.E.V. conducted the experiments.

## Notes

The authors declare no competing financial interest.

## References

1. Moore, S. J.; MacDonald, J. T.; Wienecke, S.; Ishwarbhai, A.; Tsipa, A.; Aw, R.; Kylilis, N.; Bell, D. J.; McClymont, D. W.; Jensen, K.; Polizzi, K. M.; Biedendieck, R.; Freemont, P. S., Rapid acquisition and model-based analysis of cell-free transcription-translation reactions from nonmodel bacteria. Proc Natl Acad Sci U S A 2018, 115(19), E4340–E4349.

2. Chappell, J.; Jensen, K.; Freemont, P. S., Validation of an entirely in vitro approach for rapid prototyping of DNA regulatory elements for synthetic biology. Nucleic Acids Res 2013, 41(5), 3471–81.

3. Senoussi, A.; Lee Tin Wah, J.; Shimizu, Y.; Robert, J.; Jaramillo, A.; Findeiss, S.; Axmann, I. M.; Estevez-Torres, A., Quantitative Characterization of Translational Riboregulators Using an in Vitro Transcription-Translation System. ACS Synth Biol 2018, 7 (5), 1269–1278.

4. Lehr, F. X.; Hanst, M.; Vogel, M.; Kremer, J.; Goringer, H. U.; Suess, B.; Koeppl, H., Cell-Free Prototyping of AND-Logic Gates Based on Heterogeneous RNA Activators. ACS Synth Biol 2019, 8(9), 2163–2173.

5. Wang, H.; Li, J.; Jewett, M. C., Development of a Pseudomonas putida cell-free protein synthesis platform for rapid screening of gene regulatory elements. Synthetic Biology 2018, 3(1).

6. Karim, A. S.; Jewett, M. C., A cell-free framework for rapid biosynthetic pathway prototyping and enzyme discovery. Metab Eng 2016, 36, 116–126.

7. Casini, A.; Chang, F. Y.; Eluere, R.; King, A.M.; Young, E. M.; Dudley, Q. M.; Karim, A.; Pratt, K.; Bristol, C.; Forget, A.; Ghodasara, A.; Warden-Rothman, R.; Gan, R.; Cristofaro, A.; Borujeni, A. E.; Ryu, M. H.; Li, J.; Kwon, Y. C.; Wang, H.; Tatsis, E.; Rodriguez-Lopez, C.; O’Connor, S.; Medema, M. H.; Fischbach, M. A.; Jewett, M. C.; Voigt, C.; Gordon, D. B., A Pressure Test to Make 10 Molecules in 90 Days: External Evaluation of Methods to Engineer Biology. JAm Chem Soc 2018, 140(12), 4302–4316.

8. Pardee, K.; Green, A. A.; Takahashi, M. K.; Braff, D.; Lambert, G.; Lee, J.W.; Ferrante, T.; Ma, D.; Donghia, N.; Fan, M.; Daringer, N. M.; Bosch, I.; Dudley, D. M.; O’Connor, D. H.; Gehrke, L.; Collins, J. J., Rapid, Low-Cost Detection of Zika Virus Using Programmable Biomolecular Components. Cell 2016, 165(5), 1255–1266.

9. Thavarajah, W.; Silverman, A. D.; Verosloff, M. S.; Kelley-Loughnane, N.; Jewett, M. C.; Lucks, J. B., Point-of-Use Detection of Environmental Fluoride via a Cell-Free Riboswitch-Based Biosensor. ACS Synth Biol 2020, 9(1), 10–18.

10. Alam, K. K.; Jung, J. K.; Verosloff, M. S.; Clauer, P. R.; Lee, J. W.; Capdevila, D. A.; Pastén, P. A.; Giedroc, D. P.; Collins, J. J.; Lucks, J. B., 2019.

11. Wen, K. Y.; Cameron, L.; Chappell, J.; Jensen, K.; Bell, D.J.; Kelwick, R.; Kopniczky, M.; Davies, J. C.; Filloux, A.; Freemont, P. S., A Cell-Free Biosensor for Detecting Quorum Sensing Molecules in P. aeruginosa-infected Respiratory Samples. ACS Synth Biol 2017, 6(12), 2293–2301.

12. Sun, Z.Z.; Hayes, C. A.; Shin, J.; Caschera, F.; Murray, R. M.; Noireaux, V., Protocols for implementing an Escherichia coli based TX-TL cell-free expression system for synthetic biology. J Vis Exp 2013, (79), e50762.

13. Zimmerman, S. B.; Trach, S. O., Estimation of macromolecule concentrations and excluded volume effects for the cytoplasm of Escherichia coli. J Mol Biol 1991, 222(3), 599–620.

14. Jewett, M. C.; Swartz, J. R., Mimicking the Escherichia coli cytoplasmic environment activates long-lived and efficient cell-free protein synthesis. Biotechnol Bioeng 2004, 86(1), 19–26.

15. Kuznetsova, I. M.; Zaslavsky, B. Y.; Breydo, L.; Turoverov, K. K.; Uversky, V. N., Beyond the excluded volume effects: mechanistic complexity of the crowded milieu. Molecules 2015, 20(1), 1377-409.

16. Kontur, W. S.; Capp, M. W.; Gries, T. J.; Saecker, R. M.; Record, M. T.Jr., Probing DNA binding, DNA opening, and assembly of a downstream clamp/jaw in Escherichia coli RNA polymerase-lambdaP(R) promoter complexes using salt and the physiological anion glutamate. Biochemistry 2010, 49(20), 4361-73.

17. Ge, X.; Luo, D.; Xu, J., Cell-free protein expression under macromolecular crowding conditions. PLoS One 2011, 6(12), e28707.

18. Moriizumi, Y.; Tabata, K. V.; Miyoshi, D.; Noji, H., Osmolyte-Enhanced Protein Synthesis Activity of a Reconstituted Translation System. ACS Synth Biol 2019, 8(3), 557–567.

19. Shin, J.; Noireaux, V., An E. coli cell-free expression toolbox: application to synthetic gene circuits and artificial cells. ACS Synth Biol 2012, 1(1), 29–41.

20. Espah Borujeni, A.; Channarasappa, A. S.; Salis, H. M., Translation rate is controlled by coupled trade-offs between site accessibility, selective RNA unfolding and sliding at upstream standby sites. Nucleic Acids Res 2014, 42(4), 2646–59.

21. Salis, H. M.; Mirsky, E. A.; Voigt, C. A., Automated design of synthetic ribosome binding sites to control protein expression. Nat Biotechnol 2009, 27 (10), 946–50.

22. Kai, L.; Dotsch, V.; Kaldenhoff, R.; Bernhard, F., Artificial environments for the co-translational stabilization of cell-free expressed proteins. PLoS One 2013, 8(2), e56637.

23. Karim, A. S.; Heggestad, J. T.; Crowe, S. A.; Jewett, M. C., Controlling cell-free metabolism through physiochemical perturbations. Metab Eng 2018, 45, 86–94.

24. Espah Borujeni, A.; Salis, H. M., Translation Initiation is Controlled by RNA Folding Kinetics via a Ribosome Drafting Mechanism. J Am Chem Soc 2016, 138(22), 7016–23.

25. Reis, A. C.; Salis, H. M., An Automated Model Test System for Systematic Development and Improvement of Gene Expression Models. ACS Synth Biol 2020, 9(11), 3145–3156.

26. Farasat, I.; Kushwaha, M.; Collens, J.; Easterbrook, M.; Guido, M.; Salis, H. M., Efficient search, mapping, and optimization of multi-protein genetic systems in diverse bacteria. Mol Syst Biol 2014, 10, 731.

27. Niess, A.; Siemann-Herzberg, M.; Takors, R., Protein production in Escherichia coli is guided by the trade-off between intracellular substrate availability and energy cost. Microb Cell Fact 2019, 18(1), 8.

28. Klumpp, S.; Scott, M.; Pedersen, S.; Hwa, T., Molecular crowding limits translation and cell growth. Proc Natl Acad Sci U S A 2013, 110(42), 16754–9.

29. Di̇nç, C. Ö.; ICibarer, G.; Giiner, A., Solubility profiles of poly(ethylene glycol)/solvent systems. II. comparison of thermodynamic parameters from viscosity measurements. Journal of Applied Polymer Science 2010, 117(2), 1100–1119.

30. Borg, A.; Ehrenberg, M., Determinants of the rate of mRNA translocation in bacterial protein synthesis. J Mol Biol 2015, 427(9), 1835–47.

31. Yamagami, R.; Bingaman, J. L.; Frankel, E. A.; Bevilacqua, P. C., Cellular conditions of weakly chelated magnesium ions strongly promote RNA stability and catalysis. Nat Commun 2018, 9(1), 2149.

32. Wohlgemuth, I.; Pohl, C.; Rodnina, M. V., Optimization of speed and accuracy of decoding in translation. EMBO J 2010, 29(21), 3701–9.

33. Lebowitz, J. L.; Helfand, E.; Praestgaard, E., Scaled Particle Theory of Fluid Mixtures. The Journal of Chemical Physics 1965, 43(3), 774–779.

34. Ling, K.; Jiang, H.; Zhang, Q., A colorimetric method for the molecular weight determination of polyethylene glycol using gold nanoparticles. Nanoscale Res Lett 2013, 8(1), 538.

35. Allen, S. H.; Wong, K. P., The hydrodynamic and spectroscopic properties of 16 S RNA from Escherichia coli ribosome in reconstitution buffer. J Biol Chem 1978, 253(24), 8759–66.

36. Gabler, R.; Westhead, E. W.; Ford, N. C., Studies of ribosomal diffusion coefficients using laser light-scattering spectroscopy. BiophysJ 1974, 14(7), 528–45.

37. Werner, A., Predicting translational diffusion of evolutionary conserved RNA structures by the nucleotide number. Nucleic Acids Res 2011, 39(3), e17.

38. Sullivan, C. J.; Pendleton, E. D.; Sasmor, H. H.; Hicks, W. L.; Farnum, J.B.; Muto, M.; Amendt, E. M.; Schoborg, J. A.; Martin, R.W.; Clark, L. G.; Anderson, M. J.; Choudhury, A.; Fior, R.; Lo, Y. H.; Griffey, R. H.; Chappell, S. A.; Jewett, M. C.; Mauro, V. P.; Dresios, J., A cell-free expression and purification process for rapid production of protein biologics. Biotechnol J 2016, 11(2), 238–48.

39. Stark, J. C.; Jaroentomeechai, T.; Moeller, T. D.; Dubner, R.S.; Hsu, K. J.; Stevenson, T. C.; DeLisa, M. P.; Jewett, M. C., On-demand, cell-free biomanufacturing of conjugate vaccines at the point-of-care. bioRxiv 2019, 681841.

40. Fritz, B. R.; Jamil, O. K.; Jewett, M. C., Implications of macromolecular crowding and reducing conditions for in vitro ribosome construction. Nucleic Acids Res 2015, 43(9), 4774–84.

41. Rustad, M.; Eastlund, A.; Jardine, P.; Noireaux, V., Cell-free TXTL synthesis of infectious bacteriophage T4 in a single test tube reaction. Synthetic Biology 2018, 3(1).

42. Jung, J. K.; Alam, K. K.; Verosloff, M. S.; Capdevila, D. A.; Desmau, M.; Clauer, P. R.; Lee, J. W.; Nguyen, P. Q.; Pasten, P. A.; Matiasek, S. J.; Gaillard, J. F.; Giedroc, D. P.; Collins, J. J.; Lucks, J. B., Cell-free biosensors for rapid detection of water contaminants. Nat Biotechnol 2020, 38(12), 1451–1459.

43. Takahashi, M. K.; Hayes, C. A.; Chappell, J.; Sun, Z. Z.; Murray, R. M.; Noireaux, V.; Lucks, J. B., Characterizing and prototyping genetic networks with cell-free transcription-translation reactions. Methods 2015, 86, 60–72.

44. de los Santos, E. L.; Meyerowitz, J. T.; Mayo, S. L.; Murray, R. M., Engineering Transcriptional Regulator Effector Specificity Using Computational Design and In Vitro Rapid Prototyping: Developing a Vanillin Sensor. ACS Synth Biol 2016, 5 (4), 287–95.

45. Shin, J.; Noireaux, V., Efficient cell-free expression with the endogenous E. Coli RNA polymerase and sigma factor 70. J Biol Eng 2010, 4, 8.

46. Pardee, K.; Slomovic, S.; Nguyen, P. Q.; Lee, J.W.; Donghia, N.; Burrill, D.; Ferrante, T.; McSorley, F. R.; Furuta, Y.; Vernet, A.; Lewandowski, M.; Boddy, C. N.; Joshi, N. S.; Collins, J. J., Portable, On-Demand Biomolecular Manufacturing. Cell 2016, 167 (1), 248–259 e12.

47. Tjorve, K. M. C.; Tjorve, E., A proposed family of Unified models for sigmoidal growth. Ecol Model 2017, 359, 117–127.

